# Discovery of a persistent Zika virus lineage in Bahia, Brazil

**DOI:** 10.1101/049916

**Authors:** Samia N. Naccache, Julien Thézé, Silvia I. Sardi, Sneha Somasekar, Alexander L. Greninger, Antonio C. Bandeira, Gubio S. Campos, Laura B. Tauro, Nuno R. Faria, Oliver G. Pybus, Charles Y. Chiu

**Author notes:** Co-first authors (Samia N. Naccache, Julien Thézé). Corresponding author: Charles Chiu, MD/PhD.

## Abstract

Metagenomic next-generation sequencing coupled with capture probe enrichment was used to recover 11 whole and partial Zika virus (ZIKV) genomes from patients in Bahia, Brazil from April 2015 to January 2016, where the majority of suspected Brazilian ZIKV cases have been reported. Phylogenetic reconstructions and molecular clock analyses using the newly generated data uncovered the existence of a Bahia-specific ZIKV lineage sharing a common ancestor in mid-2014, indicating sustained circulation of this strain in Bahia since that date.

## Introduction

Zika virus is an arthropod-borne RNA virus primarily transmitted by mosquitoes of the *Aedes* species, although transmission through blood transfusion and sexual contact has also been described (1). ZIKV can be classified in two genotypes: the African genotype, found only in the African continent, and the Asian genotype, associated with outbreaks in Southeast Asia, several Pacific islands and, more recently, in the Americas (2). In May of 2015, Brazil reported the first detection of autochthonous cases of ZIKV in the northeast of the country (3, 4). Since then, transmission has been confirmed in 22 of the 27 federal states (5).

The rapid geographic expansion of ZIKV transmission, together with its proposed association with microcephaly and congenital abnormalities (6), demand a rapid increase in molecular surveillance in the most affected areas. This is particularly relevant for regions where other mosquito-borne viruses, particularly dengue virus (DENV) and chikungunya virus (CHIKV), are known to co-circulate with ZIKV (2), as surveillance based on clinical symptoms alone can lead to misdiagnosed cases. Accurate genetic characterization of circulating strains can also facilitate determination of the origin and potential spread of ZIKV infection in travelers returning from endemic countries. Previous analyses have suggested that ZIKV was introduced in the Americas at least one year before its initial detection in Brazil (1). Bahia state comprised 93% of suspected ZIKV cases in Brazil in 2015 (2), and has reported cases of ZIKV-associated fetal microcephaly, although no genetic information aside from one complete genome is available from the region to date (2, 7). Here we report epidemiological findings obtained using 11 new complete and partial ZIKV genomes from 15 clinical cases seen at Hospital Aliança in Salvador, Bahia, between April 2015 and January 2016.

## Materials and Methods

Symptomatic patients were diagnosed with acute ZIKV infection by positive qualitative RT-PCR testing from serum using primers targeting the NS5 gene (8). Metagenomic next-generation sequencing (NGS) libraries were constructed following DNase treatment of nucleic acid extracts as previously described (9, 10), followed by pathogen detection using the SURPI bioinformatics pipeline (11). Clinical samples with titers exceeding 10^4^ copies per mL by ZIKV quantitative RT-PCR using the QuantiTect SYBR Green RT-PCR kit (Qiagen) and primers targeting the fiber gene (12) produced sufficient metagenomic data to permit complete ZIKV genome assembly. For the remaining samples at lower titers, metagenomic NGS libraries were enriched for ZIKV sequences using a set of 299 XGen biotinylated lockdown capture probes (IDT Technologies) designed to tile across all ZIKV genomes (>10 kB) in the National Center for Biotechnology Information (NCBI) GenBank database as of February 2016, and curated for redundancy at a 99% similarity cutoff. Enrichment was performed using the XGen lockdown protocol and SeqCap EZ Hybridization and Wash Kit (Roche Molecular Systems) according to the manufacturer’s instructions.

Using the MAFFT program, the 11 sequences from Bahia (GenBank accession numbers KU0940224, KU0940227, KU0940228, and KX101060-KX101067) were aligned together with all published and available near-complete ZIKV genomes and longer sub-genomic regions (>1500nt) of the Asian genotype as of April of 2016. Both maximum likelihood (ML) and Bayesian phylogenetic inferences were performed using PhyML and BEASTv1.8.2 programs, respectively. The ML phylogeny was reconstructed using the general time reversible nucleotide substitution model with a proportion of invariant sites (GTR+I). Statistical support for phylogenetic nodes was assessed using a bootstrap approach (with 100 replicates). A Bayesian molecular clock phylogeny was estimated using the best fitting evolutionary model determined in Faria, et al (1); specifically, a strict molecular clock GTR+I substitution model, a Bayesian skyline coalescent prior and a non-informative continuous time Markov chain (CTMC) reference prior for the molecular clock rate.

## Results

In total, 11 ZIKV partial and complete genome sequences were recovered, with average coverage of 69.4 ± 2.0% and coverage ranging from 40 - 100% (Table 1). Nearly all isolates from Bahia (11 out of 12) clustered together within a single strongly supported clade (posterior probability 1.00, bootstrap support 96%, Clade C, Figure 1). This support for the Bahia clade is notable given that the majority of ZIKV genomes are incomplete, and this uncertainty is accounted for in the phylogenetic inference. The tree topology obtained is in accordance with previous findings (1,4,6) and the time to the most recent common ancestor (TRMCA) of the epidemic in the Americas is similar to that previously estimated (1) (Clade A, Figure 1). Our analyses using the new genomes generated here indicate that ZIKV was introduced in Bahia between March and September 2014. An isolate from Maranhão, northeast Brazil, located 1000 km from Bahia, is ancestral to the Bahia clade (bootstrap support 66%, posterior probability 1.00, Clade B, Figure 1). The TRMCA of this northeast Brazil clade (comprising the Bahia clade and the Maranhão sequence) is estimated to be between September 2013 and April 2014, at a very early stage of the epidemic, strengthening the evidence for the hypothesis that ZIKV in the Americas originated from Brazil (1). One previously reported sequence from Bahia (7) clustered (posterior probability=0.99) with an isolate from Belém, Pará state in northern Brazil, which is 3000 km from Bahia. This may represent a separate introduction of ZIKV to Bahia state, as the patient had no history of traveling abroad and denied any sexual contact.

**Table 1.**
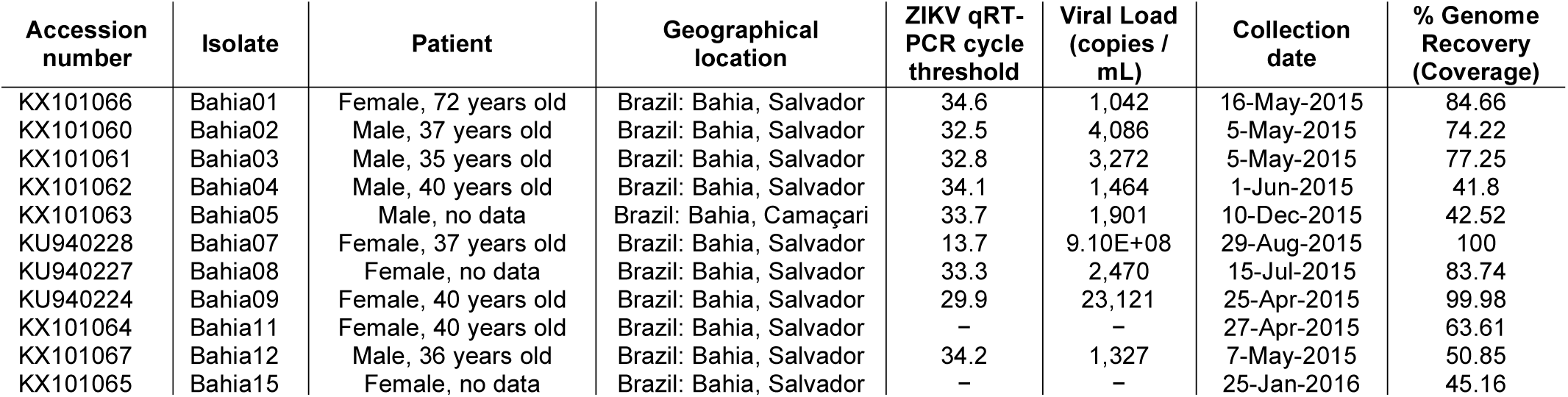
**Clinical sample information from patients with acute ZIKV infection**.

**Figure 1.**
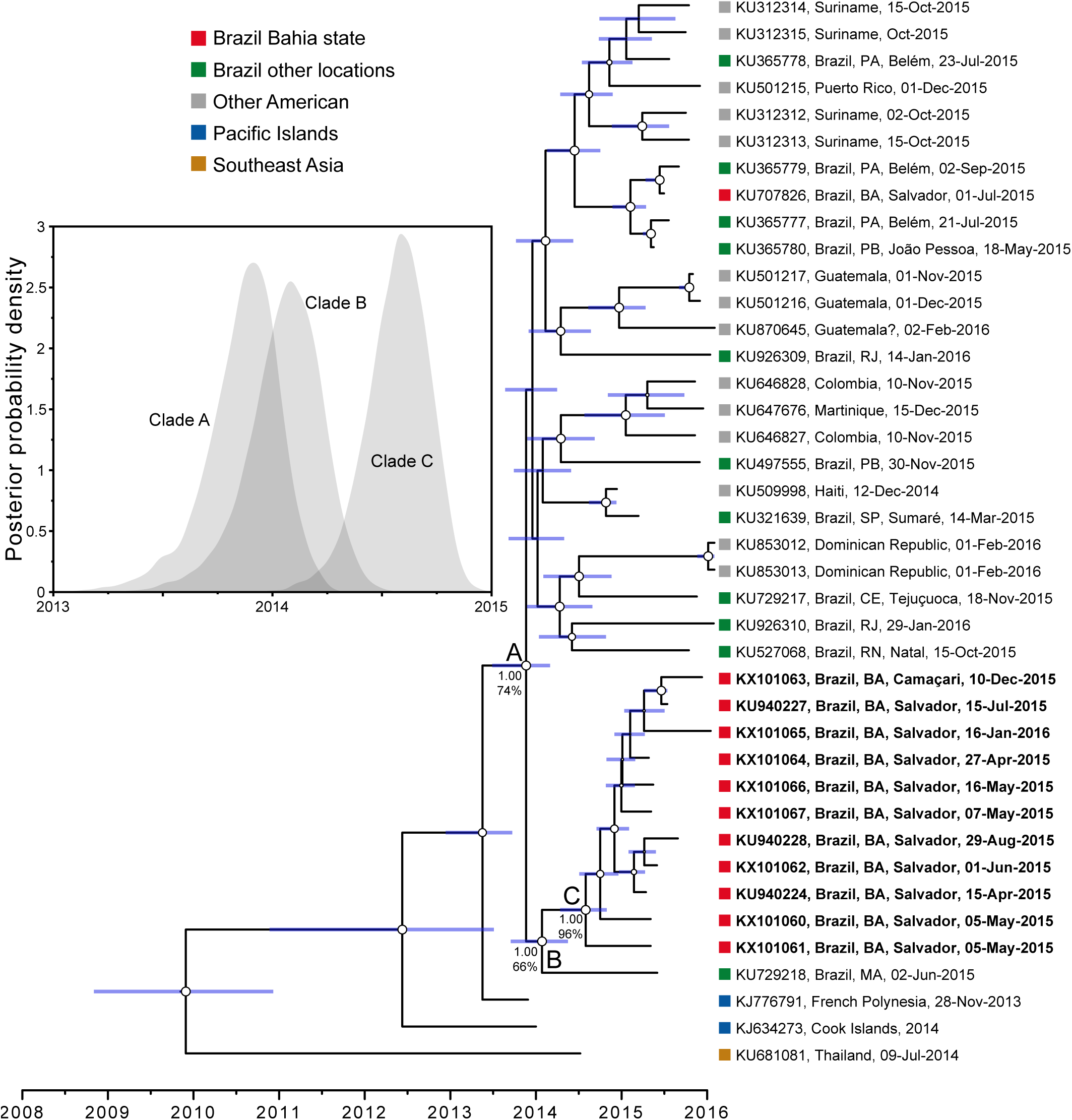
Timescale of the ZIKV outbreak in the Americas. A molecular clock phylogeny is shown of the ZIKV outbreak lineage estimated from complete and partial (>1500nt) coding region sequences. For visual clarity, five basal Southeast Asia sequences, HQ23499 (Malaysia, 1966), EU545988 (Micronesia, 2007), KU681082 (Philippines, 2012), JN860885 (Cambodia, 2010) and KU681081 (Thailand, 2013) are not displayed. Blue horizontal bars represent 95% Bayesian credible intervals for divergence dates. A, B and C denote clades discussed in main text and numbers next to them denote posterior probabilities and bootstrap scores (in %). Circle sizes represent, at each node, the posterior probability support of that node. Taxa are labeled with accession number, sampling location, and sampling date. Names of sequences generated in this study are in bold. The inset graph on the left shows the posterior distributions of the estimated ages (TMRCAs) of clades A, B and C, estimated in BEAST using the best fitting evolutionary model.

## Conclusions

Our results suggest an early introduction and presence (mid-2014) of ZIKV in the state of Bahia, Brazil. The size and statistical support for this cluster make it very likely that this lineage represents a large and sustained chain of transmission within the state. The majority of the cases of the Bahia-specific ZIKV lineage clustered closely to a sequence from Maranhão, and we found evidence for an additional autochthonous transmission event to Bahia from Pará state. Thus, ZIKV in Bahia in mid-2014 is likely to have been introduced from other regions in Brazil rather than from outside the country. The emergence of ZIKV in Bahia state in mid-2014 reported here is consistent with viral infection in pregnant women during the first trimester corresponding to the initial cases of fetal microcephaly reported in Bahia beginning in January 2015 (13), although the peak period of microcephaly did not occur until November of that year.

We cannot yet determine whether the lineage (clade C) identified here comprises the majority of ZIKV cases in Bahia, or, alternatively, whether multiple genetically distinct lineages co-circulate in the state. Brazil currently faces a major public health challenge with co-circulation of ZIKV, DENV, and CHIKV throughout the country (2-4, 14, 15). Additional molecular surveillance in the Americas and beyond is urgently needed to trace and predict the transmission of ZIKV virus.

## Acknowledgements

We thank multiple researchers worldwide for permission to include their unpublished ZIKV genomes in our analysis. This study was supported in part by Fundação de Amparo a Pesquisa do Estado da Bahia (SS), the European Research Council under the European Union’s Seventh Framework Programme (FP7/2007-2013)/ERC grant agreement no. 614725-PATHPHYLODYN (OGP), National Institutes of Health (NIH) grants R01-HL105704 and R21-AI120977 (CYC), and an award from Abbott Laboratories, Inc. (CYC). This study is made possible in part by the generous support of the American people through the United States Agency for International Development (USAID) Emerging Pandemic Threats Program - PREDICT. The contents are the responsibility of the authors and do not necessarily reflect the views of USAID or the U.S. government.

